# BioRSP: a method for characterizing enrichment patterns in single-cell embeddings

**DOI:** 10.1101/2024.06.25.599250

**Authors:** Zeyu Yao, Jake Y. Chen

**Affiliations:** Stamford American International School, Singapore 357684; Department of Biomedical Informatics and Data Science, Systems Pharmacology AI Research Center, the University of Alabama at Birmingham, Birmingham, Alabama, US

**Author notes:** Corresponding author: Dr. Jake Chen.

## Abstract

Low-dimensional embeddings such as UMAP and t-SNE are routinely used to visually interpret high-dimensional omics data, yet claims based on embedding geometry are often qualitative, embedding-sensitive, and weakly calibrated. We present BioRSP (Biological Radar Scanning Plots), a geometry-first framework that quantifies how a user-defined foreground subset is distributed across the embedding footprint of a fixed analysis set. BioRSP converts 2D coordinates to polar form around a robust vantage point, scans the set by angle, and computes a radar profile that summarizes signed radial enrichment of foreground relative to background within sliding angular windows using a distance-based radial discrepancy. The profile is reduced to interpretable summaries including anisotropy magnitude, peak directionality, and coverage, and is accompanied by explicit adequacy rules and subsampling-based stability diagnostics so the method can abstain when the geometry is underpowered. We demonstrate BioRSP in a community-standard human kidney single-nucleus reference by analyzing thick ascending limb (TAL) nuclei using published UMAP coordinates and a standardized within-set top-decile foreground rule. Within TAL, BioRSP distinguishes sharply localized rim-enrichment patterns, broadly supported but structured within-type heterogeneity, and near-isotropic profiles, with anisotropy spanning more than an order of magnitude in the pooled TAL analysis. Donor-aware reruns show that per-donor adequacy is frequently limiting, but that directional profiles are stable when donor-level support is sufficient. BioRSP is provided as open-source software producing standardized plots, summary tables, and run manifests to support reproducible embedding-aligned enrichment analysis.

## 1. Introduction

Low-dimensional neighbor embeddings such as UMAP and t-SNE are now routine for exploring single-cell and other high-dimensional omics data (McInnes et al., 2018; van der Maaten and Hinton, 2008). In these layouts, investigators commonly interpret gradients, asymmetries, and patchy structure as biological organization, including developmental progression, injury responses, tissue zonation, and condition-specific shifts (Becht et al., 2018; Farrell et al., 2018; Yost et al., 2019). Yet this interpretive step remains weakly formalized. Most quantitative approaches either partition cells into discrete groups or assess how faithfully an embedding preserves local neighborhoods. Far fewer methods directly address a common question that motivates visual inspection: whether a biologically defined foreground population is non-uniformly distributed across the footprint of an embedded cell set, and if so, where that enrichment is concentrated within the 2D layout. Polar Gini Curves partially closed this gap by quantifying how uniformly gene-positive cells occupy a cluster footprint, thereby distinguishing broadly expressed markers from patchy signals (Nguyen et al., 2021). However, as embedding-centric analyses become more widely used, geometry-grounded claims are expected to be accompanied by explicit diagnostics for artifact susceptibility, sampling limitations, and reproducibility rather than relying on qualitative inspection alone (Kobak and Berens, 2019; Gorin et al., 2022; Chari and Pachter, 2023; Johnson et al., 2022; Saquib et al., 2022; Hart and Tavolara, 2024).

Here, we introduce BioRSP (Biological Radar Scanning Plots), a geometry-first framework for quantifying angularly resolved radial enrichment of a foreground population in a two-dimensional representation. BioRSP operates on a user-defined analysis set (e.g., an annotated cell type or subtype) and a user-defined foreground rule (e.g., cells exceeding a feature-specific threshold or cells belonging to a condition of interest). Given fixed 2D coordinates, BioRSP selects a robust vantage point, transforms coordinates into polar form, and scans the analysis set by angle using sliding windows. Within each angular window, BioRSP compares the radial distributions of foreground and background cells using a signed, distance-based radial discrepancy, yielding a radar profile R(\theta) over the full angular range. We then reduce R(\theta) to a compact set of interpretable summaries, including an anisotropy magnitude that captures spatial structure strength, peak angles that localize extremal enrichment, and a coverage measure that reports how much of the profile is supported by adequate data.

BioRSP is neither a clustering nor a trajectory method. Instead, it treats published labels and user-defined subsets as strata for analysis and provides a standardized way to describe how a foreground is arranged within a given embedding footprint. Because embedding-aligned statistics can be underpowered or unstable when foreground support is sparse or uneven across angles, BioRSP includes explicit adequacy rules that mask under-supported angular sectors and can abstain from reporting scalar summaries when geometry is not identifiable. To support reproducible use, BioRSP additionally provides subsampling-based stability diagnostics that quantify the reproducibility of radar profiles and anisotropy summaries under controlled resampling, and it records analysis parameters and random seeds in machine-readable output manifests.

In this manuscript, we detail the BioRSP estimand, summary statistics, adequacy rules, and robustness diagnostics, and we demonstrate the method in a community-standard human kidney single-nucleus reference by analyzing thick ascending limb (TAL) nuclei using published UMAP coordinates and a standardized within-set top-decile foreground rule. We further perform donor-aware reruns to illustrate how per-donor sample size and foreground support limit inference, and when donor-level profiles are stable once adequacy is met. Together, these components position BioRSP as a practical, embedding-aligned framework for characterizing enrichment patterns within fixed analysis sets while explicitly reporting when the available geometry does not support reliable interpretation.

## 2. Method

### 2.1. Inputs and notation

We analyze a dataset of *N* cells indexed by *i* ∈ {1, …,*N*} Each cell *i* has a two-dimensional coordinate *z*_*i*_ ∈ ℝ^2^, representing a user-supplied embedding or 2D layout (e.g., UMAP, t-SNE, or a custom projection). We do not assume Euclidean distances in this embedding are globally faithful. Instead, the embedding is treated as a coordinate system for defining angular directions and radial distances relative to a reference point.

For each feature *g* (e.g., a gene, protein abundance, chromatin accessibility score, module score, or pathway activity score), let 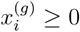 denote the feature value in cell *i* after user-chosen preprocessing/normalization. All definitions apply unchanged provided 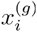 is numeric and comparable within the analyzed cell set.

BioRSP operates on a user-defined cell set *S* ⊆ {1, …,*N*} provided upstream (e.g., by filtering the input matrix and coordinates to the desired subset). The current implementation does not perform clustering or metadata-based subsetting; all quantities are computed on the provided cells.

When available, per-cell library size (total counts) is denoted by *u*_*i*_. In the current implementation, *u*_*i*_ is used only for optional stratified permutation inference; it does not affect the radar estimand itself. If *u*_*i*_ is not provided, the CLI may fall back to using per-cell expression row-sums as a proxy, which is recommended only when those row-sums reflect true library sizes.

### 2.2 Vantage point and polar coordinates

A vantage point *v* ∈ ℝ^2^ serves as the origin for radar scanning. By default, *v* is set to the geometric median of {*z*_*i*_: *i* ∈ *S*}, which is robust to outliers and irregular embedding boundaries. The geometric median is computed via an iterative procedure with fixed convergence tolerance recorded in the run manifest.

For each cell *i* ∈ *S*, we define polar coordinates relative to *v*:

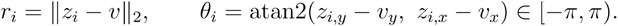

Let Θ = {*θ*^(1)^,…, *θ* ^(*B*)^} be an equally spaced grid on [− *π, π* ).Unless stated otherwise, we use *B* = 360. For each grid angle *θ*, ∈ Θ, we define a sliding angular window (sector) of width Δ radians:

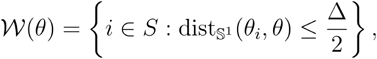

where wrapped angular distance is

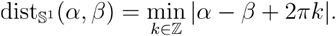

All computations use radians internally; degrees are used only for reporting. The default window width is Δ = 180^°^.

### 2.3 Foreground definition

BioRSP compares the spatial distribution of “foreground” cells (high feature values) against “background” cells (the remainder). The current implementation uses a quantile-based binary foreground to stabilize estimation across features. For feature *g* within cell set *S*, we define

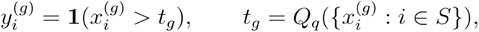

where *Q*_*q*_ is the -quantile (default *q* = 0.90) and **1**(·) is the indicator function. We use a strict inequality 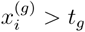 to reduce tie-related instabilities common in sparse single-cell data.

We report the realized foreground fraction

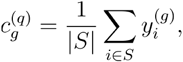

which should be interpreted as the fraction of cells assigned to the upper tail under the chosen quantile rule (not the fraction of expressing cells). If 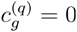 or if the total number of foreground cells is too small (adequacy rules below), the feature is treated as underpowered for stable directional estimation.

### 2.4. Radar Scanning Plot estimand

For a fixed feature *g*, cells in *S* are partitioned into a foreground set

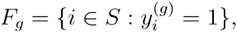

and a background set *B*_*g*_ = *S*\*F*_*g*_.

Within each angular window 𝒲 (*θ*), we compare radial samples

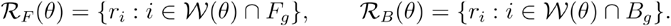

Let *W*_1_ (ℛ _*F*_, ℛ _*B*_) denote the one-dimensional Wasserstein-1 (earth mover’s) distance between the two empirical samples of radii (computed directly from the samples without binning).

BioRSP defines the signed, IQR-normalized radial discrepancy

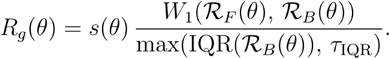

Here IQR is the interquartile range, and *τ*_IQR_ is a global IQR floor used to stabilize sectors whose background radial dispersion is near-degenerate. In the implementation,

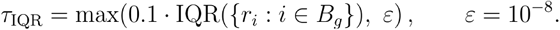

The sign term is

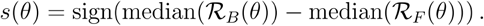

With this convention, *R*_*g*_(*θ*) > 0 indicates that in direction, foreground cells are more proximal (smaller radii; closer to the vantage point) than background cells, while *R*_*g*_ (*θ*) < 0 indicates that foreground cells are more distal (larger radii; rim-enriched) than background cells. The software records whether *τ*_IQR_ was applied in each sector as a diagnostic.

The function *R*_*g*_ (*θ*) is evaluated on the grid Θ. Any optional smoothing is used for visualization only; all summaries and inference are computed from unsmoothed values.

### 2.5. Power and adequacy rules

Directional statistics are unreliable when computed from too few foreground or background cells. We therefore impose explicit adequacy criteria and treat under-supported sectors as missing. For each direction *θ*, define:

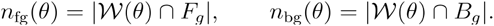

Adequacy counts are computed using the same sliding window 𝒲 (*θ*) used by the radar statistic.

A sector is considered adequate only if

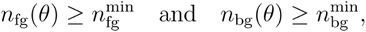

with defaults 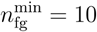 and 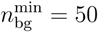. If a sector fails these criteria, *R*_*g*_ (*θ*) is set to missing and excluded from scalar summaries.

In addition to per-sector thresholds, we apply a gene-level “coverage of adequate sectors” criterion. Let

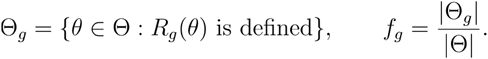

A feature is treated as adequate for downstream summarization and inference only if *f*_*g*_ ≥ *f*_min_ (default *f*_min_ = 0.9). We report *f*_*g*_ for every feature.

Finally, gene-level inference requires a minimum total number of foreground cells:

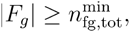

with default 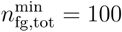. Features failing this criterion are labeled underpowered and excluded from inference (they may still be visualized cautiously).

### 2.6. Scalar summaries

Each feature’s radar profile is summarized using a primary anisotropy magnitude and interpretable peak directions computed over Θ_*g*_.

1. **Quantile foreground fraction (reported)**.

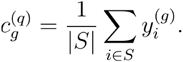
2. **Anisotropy magnitude (primary score)**. Using only valid sectors Θ_*g*_, the anisotropy score is

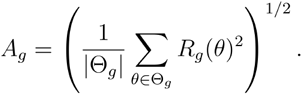

If | Θ_*g*_| *=* 0, *A*_*g*_ is reported as missing.
3. **Peak directions and strengths**. Peaks are computed over Θ_*g*_:
  a. Peak distal (rim-enriched):

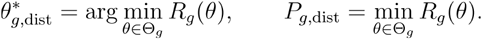
  b. Peak proximal (core-enriched):

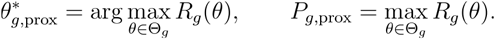
  c. Peak extremal (largest absolute magnitude):

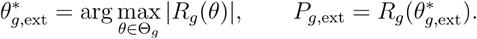

We also report 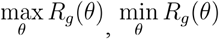, and the integrated profile as 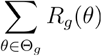 diagnostics.

### 2.7. Feature typing into four classes

To support biomarker discovery and prioritization, BioRSP can subtype features into four qualitative categories based on how widespread the feature is (coverage) and how spatially structured it is (anisotropy). Because the radar statistic uses a quantile foreground by default (which 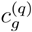 compresses across features), we recommend defining two coverage notions:

1. **Radar foreground fraction** 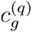: the quantile-based fraction used to compute (reported by the software).
2. **Prevalence coverage** 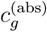: a biologically interpretable prevalence metric computed from the raw feature values, e.g.

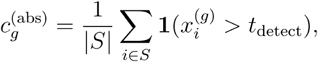

Where *t*_detect_ can be 0, a data-driven detection threshold, or a user-specified cutoff appropriate to the modality. This metric is recommended for “coverage typing” even when the radar uses quantile foreground internally.

Given a coverage metric *c*_*g*_ (typically 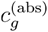 for interpretation) and anisotropy *A*_*g*_, we define thresholds *f*_hi_ and *A*_hi_ (by default, median splits across adequately powered features within S, or user-defined cutoffs recorded in the manifest). Features are then assigned to one of four types:

1. **Type I: High coverage, high anisotropy (***c*_*g*_ ≥ *c*_hi_ **and** *A*_*g*_ ≥ *A*_hi_). Widespread but strongly structured; often marks broad programs with spatial/niche bias.
2. **Type II: Low coverage, high anisotropy (***c*_*g*_ < *c*_hi_ **and** *A*_*g*_ ≥ *A*_hi_). Rare but strongly structured; high-priority candidate biomarkers.
3. **Type III: High coverage, low anisotropy (***c*_*g*_ ≥ *c*_hi_ **and** *A*_*g*_ < *A*_hi_). Broad and relatively uniform; often housekeeping-like or globally expressed programs.
4. **Type IV: Low coverage, low anisotropy (***c*_*g*_ < *c*_hi_ **and** *A*_*g*_ < *A*_hi_). Rare and weakly structured; typically lower priority unless supported by other evidence.

In all cases, features must satisfy adequacy criteria (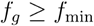 and 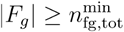) to be typed and compared reliably.

### 2.8 Pairwise feature relationships using radar profiles

To identify complementary or synergistic feature pairs, we compare their directional signatures *R*_*g*_ (*θ*) over shared valid directions. For two features *g* and *h*, define a common valid set Θ_*g,h*_ = Θ_*g*_ ⋂ Θ_*h*_.We recommend two complementary measures:

1. **Directional similarity (synergy):** correlation between radar profiles corr (*R*_*g*_ (Θ_*g,h*_), *R*_*h*_ (Θ_*g,h*_)). Strong positive correlation suggests co-localization of directional bias.
2. **Directional complementarity:** correlation between *R*_*g*_ and − *R*_*h*_, or angular separation between extremal peaks 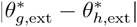 (wrapped), to detect opposing spatial biases (e.g., one distal where the other is proximal).

These pairwise analyses are performed downstream using the computed radar profiles and are recommended only for adequately powered features with stable robustness diagnostics.

### 2.9. Inference (optional permutation test)

BioRSP can assess whether the observed anisotropy exceeds what is expected if foreground membership is unrelated to spatial position in the embedding. Inference is performed on the anisotropy statistic *A*_*g*_ using permutation tests.

#### UMI-stratified permutation of foreground labels

For each feature *g*, the embedding coordinates {*z*_*i*_} (and thus {*r*_*i, θi*_} ) are held fixed. Cells are optionally partitioned into *Q* = 10 strata based on library size *u*_*i*_ (deciles within *S*), and the foreground labels 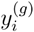 are permuted independently within each stratum. If is not provided, all cells are treated as a single stratum.

#### Fixed-direction RMS for permutations

Let 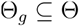 denote the set of directions where the observed radar profile is defined. To compute permutation statistics over a consistent set of directions, BioRSP uses the observed valid-direction mask Θ_*i*_ for all permutations. For permutation $k$, we compute a permuted radar profile 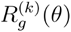 and define

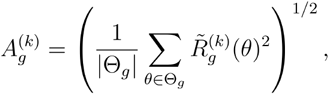

where 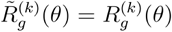 if defined and 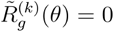 if the permuted value is missing at a direction in Θ_*g*_ If ∣ Θ_*g*_∣= 0, the p-value is reported as missing.

The empirical one-sided p-value is

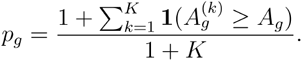

By default, *K* = 200 permutations are used for exploratory runs (and can be increased for final reporting). False discovery rate control (e.g., Benjamini–Hochberg) can be applied downstream across the set of adequately powered features.

### 2.10. Robustness diagnostics (subsampling stability)

To assess sensitivity to sampling variation, we perform repeated subsampling of cells. For each feature, we draw *M* = 20 random subsamples (default 80% of cells without replacement), recompute the radar profile and scalar summaries, and quantify two stability metrics.

Profile reproducibility is the mean Pearson correlation between the subsampled and full radar profiles, computed over directions where both profiles are finite. If fewer than two directions are jointly defined, the correlation for that subsample is recorded as missing and excluded from the mean.

Anisotropy stability is the coefficient of variation of anisotropy across subsamples:

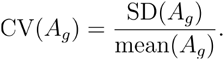

These diagnostics are reported alongside the primary summaries and are used to flag features whose directional signatures are unstable.

### 2.11. Default constants

Unless otherwise specified:

- Angular grid: *B* = 360.
- Sector width: Δ = 180 °.
- Minimum per-sector support: 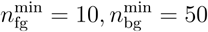.
- Minimum adequate-sector fraction: *f*_min_ = 0.9.
- Minimum total foreground for inference: 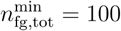.
- UMI strata: *Q* = 10.
- Numerical stabilization: ε= 10^−8^
- Permutations: *K* = 200 (exploratory; user-adjustable).
- Robustness subsampling: *M* = 20, subsample fraction =0.8.

### 2.12. Runtime complexity and scaling

Let *n* = ∣*S*∣ and *G* be the number of analyzed features. Polar coordinate computation is *O*(*n*). For each feature, the current implementation evaluates each of *B* the window centers by computing wrapped angular distances against all cells, yielding *O*(*Bn*) time to form sector masks and counts. Sector-wise Wasserstein distances are computed on radii within each window; overall runtime is dominated by the *O*(*Bn*) masking step. Permutation inference scales as *O*(*Bn*) per feature in the worst case, with underpowered features filtered by adequacy criteria prior to inference.

### 2.13. Determinism and reproducibility

All stochastic components (permutations and subsampling) are controlled by explicit random seeds recorded in output metadata. Each analysis produces machine-readable results (including adequacy diagnostics and summary statistics) and a run manifest containing parameters and seeds to support exact reproduction.

## 3. Case Study: Within–cell-type enrichment structure in the human kidney

We first demonstrate how BioRSP can be used to characterize spatial heterogeneity within a single annotated cell type in a large, community-standard single-nucleus reference dataset. We demonstrate that BioRSP complements existing marker-based analyses by systematically characterizing the distribution of high-expression cells across the embedding footprint of a fixed cell population.

We applied BioRSP to thick ascending limb (TAL) nuclei from the KPMP Human Kidney reference using the published 2D UMAP coordinates, normalized expression, TAL annotations, and donor identifiers, without reclustering or re-embedding. Using the default within-set quantile foreground rule (90th percentile; strict “>” threshold), the typical foreground size was ∼3,839 nuclei (≈10% of the TAL set; total TAL nuclei *N* ≈ 38390 inferred from the realized foreground counts).

Within the TAL cell set, we apply BioRSP gene by gene using a standardized foreground definition based on within-set expression quantiles. We first include a small panel of canonical TAL markers to serve as baseline controls, ensuring that broadly expressed markers exhibit low directional structure. We then apply BioRSP to a larger discovery panel of genes expressed in TAL cells, rank genes by the primary BioRSP anisotropy score, and retain only those genes that meet explicit adequacy and coverage criteria. This ranking step is used solely to identify representative examples of different within–cell-type enrichment phenotypes.

For statistical calibration, we compute permutation-based significance for selected genes using depth-aware shuffling within the TAL cell set, and we report corrected significance values alongside BioRSP summary metrics. To assess robustness, we repeat the analysis under controlled perturbations, including subsampling of TAL nuclei and donor-aware reruns where donor sample sizes permit.

## 4. Results

### 4.1. BioRSP reveals a spectrum of within-TAL enrichment phenotypes

Across the pooled TAL set, BioRSP anisotropy scores spanned more than an order of magnitude (*A*_*g*_ range: 0.21-2.66), demonstrating that even within a single annotated cell type, “high-expression” cells for different genes can exhibit markedly different spatial structure within the embedding footprint.

### Highly anisotropic, low-coverage focal pattern (candidate rare/focal marker behavior)

The strongest structure in this panel was observed for VCAM1 ( *A*_*g*_ = 2.66 ), which also showed substantially reduced realized coverage under the quantile foreground definition 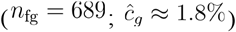, consistent with a sparse foreground caused by strong zero inflation/ties at low expression. VCAM1 exhibited an extreme directional excursion in the pooled TAL radar curve (most negative sector value *P*_*g*_ = − 5.64), indicating a sharply localized rim-enrichment (distal) phenotype in a specific angular region of the TAL embedding.

### Moderate anisotropy with typical coverage (structured within–cell-type heterogeneity)

Several genes with near-standard foreground sizes (ĉ_*g*_ ≈ 10% ) nonetheless showed clear directional structure, including TPT1 ( *A*_*g*_ = 0.93, *P*_*g*_ = − 1.91), ITGB8 ( *A*_*g*_ = 0.76, *P*_*g*_ = − 1.79), PROM1 ( *A*_*g*_ = 0.62, *P*_*g*_ = − 1.63), CDH1 (*A*_*g*_ = 0.57, *P*_*g*_ = − 1.43 ), and SLC12A1 (*A*_*g*_ = 0.32, *P*_*g*_ = − 0.69). In practical terms, these profiles indicate that the top-expression decile for these genes is not uniformly distributed over the TAL embedding footprint; instead, high-expression nuclei preferentially occupy particular angular sectors and (depending on sign) tend toward either the core or rim in those directions. This phenotype is the regime BioRSP is designed to capture for within– cell-type substructure discovery.

### Core-enriched signatures with weak-to-moderate directionality (foreground generally more proximal)

A subset of genes showed radar profiles whose minima were positive, meaning the foreground was radially more proximal than background across essentially all directions in the pooled TAL set under the current sign convention. Notably, DEFB1 ( *A*_*g*_ = 0.89, *P*_*g*_ = 0.43) and UMOD ( *A*_*g*_ = 0.81, *P*_*g*_ = 0.27) fell in this category, consistent with a predominantly “core-enriched” phenotype rather than a rim-localized one.

### Near-isotropic controls / weak structure

At the low end of the score distribution, MT-CO1 ( *A*_*g*_ = 0.21, *P*_*g*_ = − 0.61), CD24 ( *A*_*g*_ = 0.26, *P*_*g*_ = − 0.58), and ATP1A1 ( *A*_*g*_ = 0.29, *P*_*g*_ = − 0.74) showed comparatively small anisotropy magnitudes, producing flatter radar curves consistent with weak directional structure under the top-decile foreground definition.

Together, these results illustrate that BioRSP can separate (i) sparse, highly localized patterns (VCAM1-like), (ii) broadly supported but directionally structured within-type patterns (PROM1/CDH1/ITGB8-like), and (iii) comparatively uniform patterns (MT-CO1/CD24/ATP1A1-like), all while operating strictly within a fixed TAL cell set and without relying on reclustering or trajectory inference.

### Permutation calibration on selected genes

All reported genes achieved the minimum resolvable one-sided p-value under the stored permutation setting. This is consistent with evaluating a deliberately selected set enriched for pronounced structure and indicates that, at this permutation resolution, the observed anisotropy exceeded all permuted values for each tested gene.

### 4.2. Donor-aware robustness highlights power limits per donor, but stable profiles when adequate

To assess robustness to donor structure, we reran BioRSP within each donor where TAL nuclei were available (56 donors total) for a representative set of 12 genes. A key practical observation was that donor-level adequacy is often the limiting factor: for most genes, only ∼7-10 of 56 donors (≈12-18%) met the gene-level adequacy criteria under the default thresholds. VCAM1 was not adequate in any single-donor run, consistent with its sparse foreground even in the pooled TAL set.

Importantly, when a donor passed adequacy, the directional profiles were typically well-supported across angles (mean adequacy fraction ≈ 0.997 among adequate donor runs; minimum observed ≈ 0.903), indicating that failures were primarily due to insufficient TAL counts/foreground counts rather than localized angular dropout. Directional agreement across adequate donors was gene-dependent: some genes showed moderate-to-strong angular concentration (e.g., SPP1 with concentration *R* ≈ 0.68, implying relatively consistent peak direction), whereas others were more weakly aligned (e.g., UMOD, *R* ≈ 0.26), reflecting either genuine donor variability or residual sampling noise at donor-specific cell counts. Overall, these donor-aware diagnostics support the pooled TAL analysis as the primary, well-powered characterization, while also showing that donor-stratified confirmation is feasible but requires sufficient per-donor TAL nuclei (or relaxed thresholds / larger sector widths) to be informative.

## 5. Conclusion

BioRSP provides a practical, embedding-aligned statistic for quantifying enrichment structure within a fixed analysis set without reclustering, trajectory inference, or assumptions that Euclidean distances are globally meaningful in the embedding. By scanning a cell set in angular windows around a robust vantage point, BioRSP summarizes foreground-versus-background radial differences as a radar profile *R*(*θ*), and reduces that profile to interpretable outputs (anisotropy magnitude, peak directions, and coverage) accompanied by explicit adequacy and instability flags. This design directly targets a common gap in embedding-based workflows: investigators often observe gradients, asymmetries, and patchiness within an annotated population but lack a calibrated, standardized way to characterize these patterns feature-by-feature while transparently reporting when the geometry is not identifiable.

In a human kidney reference application focused on thick ascending limb (TAL) nuclei, BioRSP reveals that “high-expression” cells can occupy markedly different spatial configurations even within a single annotated cell type. Using a standardized within-set top-decile foreground rule on published UMAP coordinates, BioRSP separates sparse, highly localized rim-enrichment phenotypes (VCAM1-like) from broadly supported but directionally structured within-type patterns (PROM1/CDH1/ITGB8/SLC12A1-like), and from comparatively uniform, near-isotropic profiles (MT-CO1/CD24/ATP1A1-like). These results illustrate how BioRSP complements conventional marker-based summaries by making within-type heterogeneity explicit and comparable across features through a common estimand and reporting scheme.

Donor-aware reruns further clarify how BioRSP should be used in multi-donor atlases: donor-level adequacy is often the limiting factor, and many genes do not meet per-donor support thresholds under default settings, especially when the foreground is sparse. However, when donor runs are adequately powered, radar profiles are generally well-supported across angles, enabling donor-stratified confirmation of pooled signals in settings where sufficient per-donor cell counts exist. Practically, this indicates that pooled analyses are typically the primary, well-powered characterization, while donor-aware analyses are most informative as a robustness layer (and may require relaxed thresholds or larger sector widths when per-donor counts are modest).

BioRSP has clear boundaries that should guide interpretation. It does not assert that embedding geometry is a faithful physical space, nor does it infer developmental direction, temporal movement, or causal trajectories; instead, it provides a calibrated description of foreground distribution in the provided coordinate system. The default quantile-based foreground stabilizes comparisons across features, but prevalence-oriented interpretations should incorporate a separate, biologically meaningful detection/coverage metric when appropriate. Going forward, the same framework can be extended with additional foreground definitions (e.g., continuous weighting or covariate-adjusted thresholds), alternative coordinate systems, and richer cross-replicate models, while preserving the core principle that embedding-aligned claims should be accompanied by adequacy reporting and stability diagnostics rather than qualitative inspection alone.

## 6. Code and data availability

BioRSP is released as open-source software and is publicly available on GitHub at https://github.com/cytronicoder/biorsp. All analyses reported in this manuscript were executed from a fixed repository snapshot to ensure reproducibility.

The TAL case study uses the Azimuth Human Kidney reference, a KPMP/HuBMAP-derived single-nucleus RNA-seq reference distributed by the HuBMAP consortium. The reference is publicly available via Zenodo (DOI: 10.5281/zenodo.10694842). For analysis, the reference was converted from the distributed Seurat object format into an AnnData (.h5ad) representation using utilities provided in the BioRSP repository. The exact reference file used in this study, along with its checksum, is documented in the accompanying run manifest.

All BioRSP analyses generate a machine-readable run manifest that records the complete computational provenance of each execution. This manifest includes:

- Analysis parameters and foreground definitions
- Random seeds and execution timestamps
- Dataset sizes and gene filtering outcomes
- Donor composition and adequacy diagnostics (where applicable)
- Paths to all generated result tables and figures

The manifest enables exact reproduction of reported results and is intended to accompany any redistributed outputs. For the TAL case study, BioRSP produces standardized result artifacts, including ranked gene summary tables, angular enrichment statistics, and visualization outputs. These files, together with the run manifest and repository commit identifier, fully specify the analysis performed in this manuscript.

## 7. Author contributions

Z.Y. conceived the study, developed the methodological framework, implemented the software, curated and processed the datasets, performed the computational analyses, and drafted the manuscript. J.Y.C. designed the experimental and evaluation framework, supervised the project, and provided scientific guidance throughout. Both authors reviewed, edited, and approved the final manuscript.

