## Supplemental File 1 for "BioRSP: a method for characterizing enrichment patterns in single-cell embeddings"

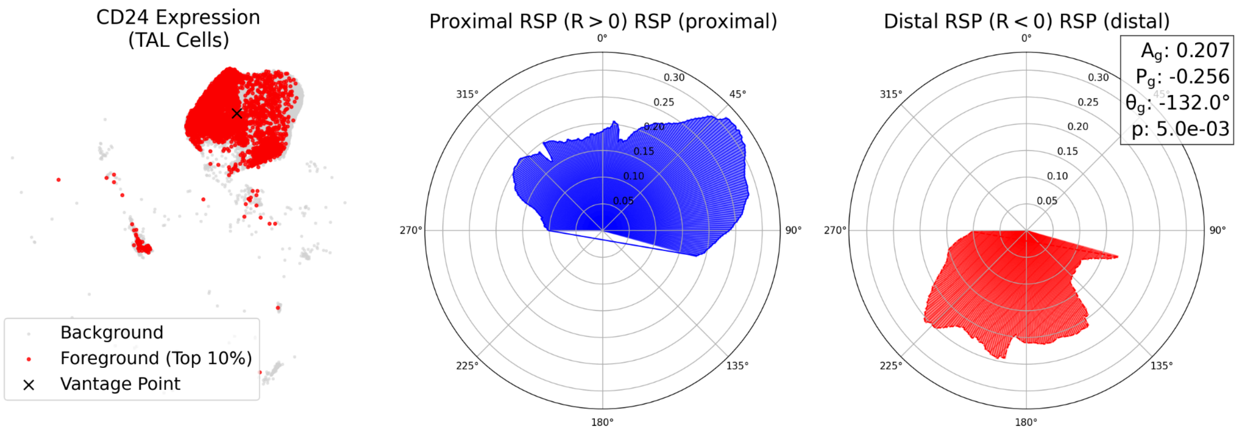


**Figure S1.** CD24 spatial enrichment profile in TAL embedding. Left: UMAP of TAL cells with CD24 foreground defined as the top 10 percent of expression (red) over background TAL cells (gray); the black x marks the BioRSP vantage point. Middle: proximal radar profile showing angles with positive radial enrichment (R greater than 0). Right: distal radar profile showing angles with negative radial enrichment (R less than 0), with summary statistics reported in the inset (Ag 0.207, Pg -0.256, theta -132.0 degrees; stratified permutation p 5.0e-03).

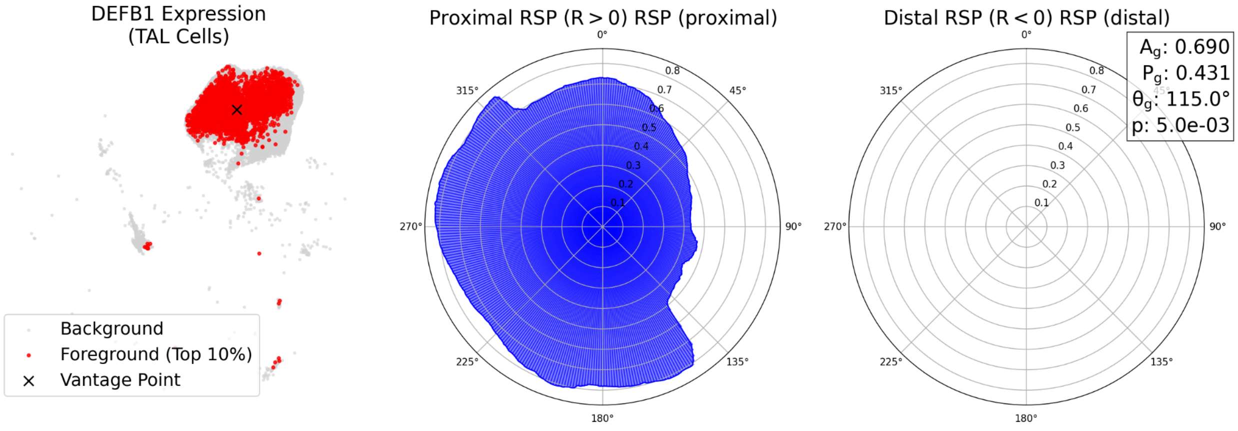


**Figure S2.** DEFB1 spatial enrichment profile in TAL embedding. Left: TAL UMAP with DEFB1 top 10 percent foreground (red), background (gray), and vantage point (x). Middle: proximal (R greater than 0) radar profile. Right: distal (R less than 0) radar profile with inset summary (Ag 0.690, Pg 0.431, theta 115.0 degrees; stratified permutation p 5.0e-03).

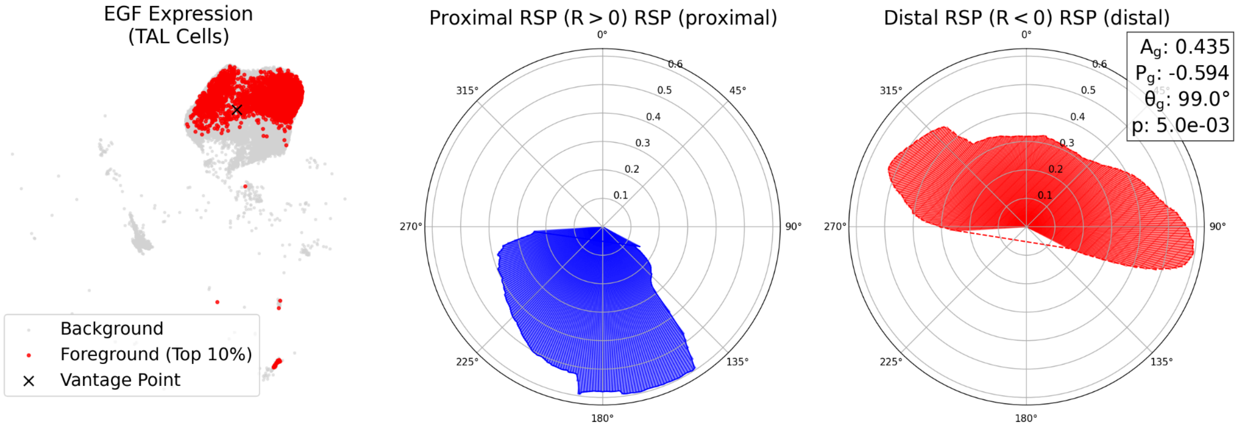


**Figure S3.** EGF spatial enrichment profile in TAL embedding. Left: TAL UMAP showing EGF top 10 percent foreground (red), background (gray), and vantage point (x). Middle: proximal (R greater than 0) enrichment profile. Right: distal (R less than 0) depletion profile with summary statistics (Ag 0.435, Pg -0.594, theta 99.0 degrees; stratified permutation p 5.0e-03).

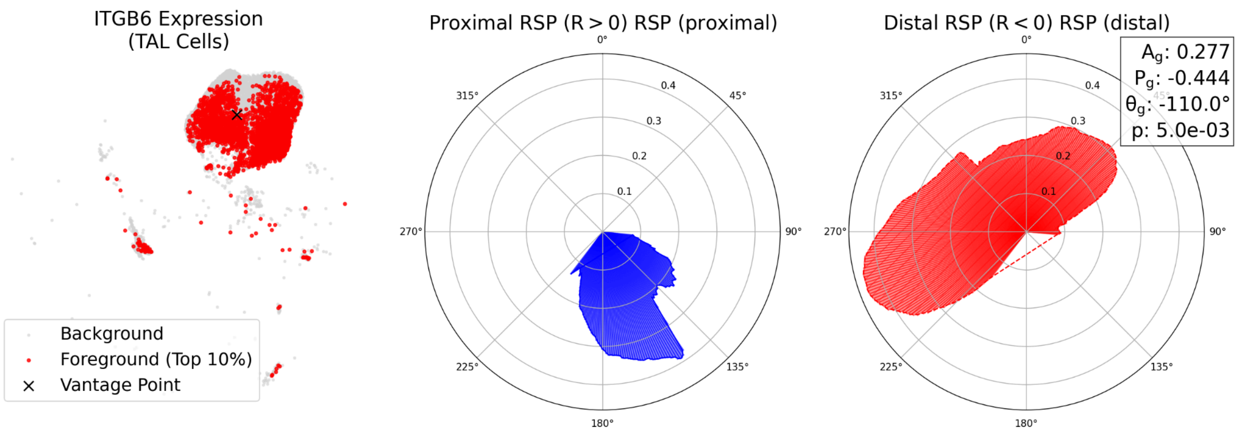


**Figure S4.** ITGB6 spatial enrichment profile in TAL embedding. Left: TAL UMAP with ITGB6 top 10 percent foreground (red), background (gray), and vantage point (x). Middle: proximal (R greater than 0) radar profile. Right: distal (R less than 0) radar profile with inset summary (Ag 0.277, Pg -0.444, theta -110.0 degrees; stratified permutation p 5.0e-03).

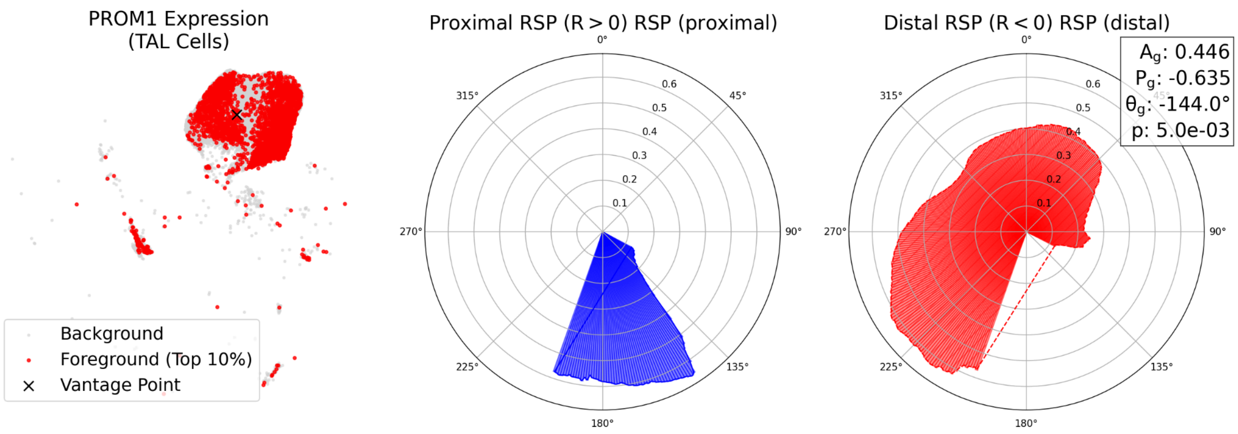


**Figure S5.** PROM1 spatial enrichment profile in TAL embedding. Left: TAL UMAP with PROM1 top 10 percent foreground (red), background (gray), and vantage point (x). Middle: proximal (R greater than 0) radar profile. Right: distal (R less than 0) radar profile with inset summary (Ag 0.446, Pg -0.635, theta -144.0 degrees; stratified permutation p 5.0e-03).

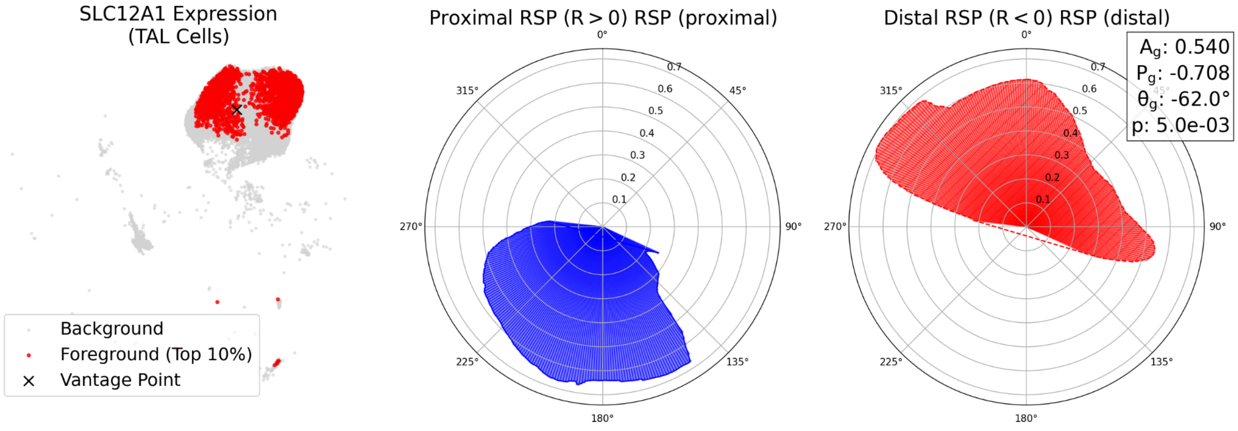


**Figure S6.** SLC12A1 spatial enrichment profile in TAL embedding. Left: TAL UMAP with SLC12A1 top 10 percent foreground (red), background (gray), and vantage point (x). Middle: proximal (R greater than 0) radar profile. Right: distal (R less than 0) radar profile with inset summary (Ag 0.540, Pg -0.708, theta -62.0 degrees; stratified permutation p 5.0e-03).

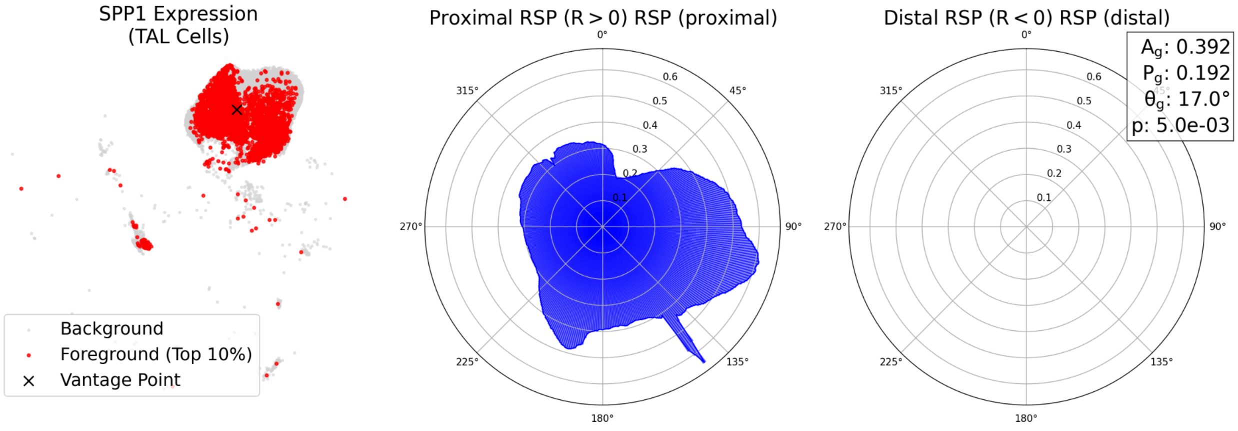


**Figure S7.** SPP1 spatial enrichment profile in TAL embedding. Left: TAL UMAP with SPP1 foreground (top 10% expression; red), background (gray), and the BioRSP vantage point (x). Middle: proximal radar profile ($R > 0$). Right: distal radar profile ($R < 0$) with inset metrics (Ag 0.392, Pg 0.192, theta 17.0 degrees; p 5.0e-03).

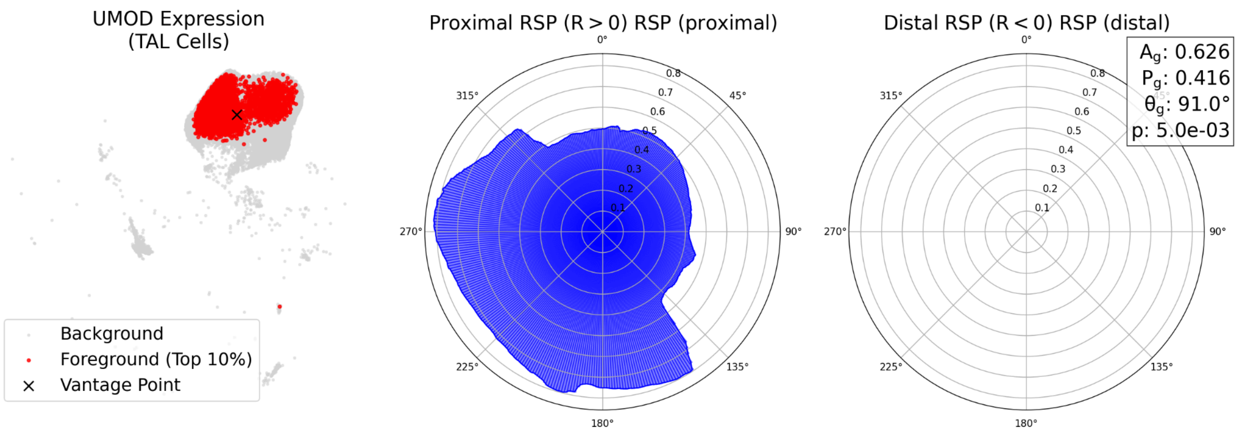


**Figure S8.** UMOD spatial enrichment profile in TAL embedding. Left: TAL UMAP with UMOD foreground (top 10% expression; red), background (gray), and the BioRSP vantage point (x). Middle: proximal radar profile (R > 0). Right: distal radar profile (R < 0) with inset metrics (Ag 0.626, Pg 0.416, theta 91.0 degrees; p 5.0e-03).
